# High-dimensional spectral cytometry panels for whole blood immune phenotyping

**DOI:** 10.1101/2023.07.17.549272

**Authors:** Tom Dott, Slobodan Culina, Rene Chemali, Cedric Ait Mansour, Florian Dubois, Bernd Jagla, Jean Marc Doisne, Lars Rogge, François Huetz, Friederike Jönsson, Pierre-Henri Commere, James Di Santo, Benjamin Terrier, Lluis Quintana-Murci, Darragh Duffy, Milena Hasan, Milieu Intérieur Consortium

**Author notes:** These authors contributed equally (Tom Dott, Slobodan Culina).

## Abstract

The need to understand the mechanisms and pathways of immune responses in pathogenic conditions such as cancer and autoimmunity requires awareness of natural immune variability in healthy subjects. To this end, various systems immunology studies have been established. Among them, the *Milieu Intérieur* (MI) study was established to define the boundaries of a healthy immune response and identify determinants of immune response variation. MI used immunophenotyping of a 1000 healthy donor cohort by flow cytometry as a principal outcome for immune variance at steady state. For the 10-year longitudinal MI study, we have developed two high-dimensional spectral flow cytometry panels that allow deep characterization of innate and adaptive whole blood immune cells (35 and 34 fluorescent markers, respectively) and standardized the protocol for sample handling, staining, acquisition, and data analysis. This permits the reproducible quantification of over 182 immune cell phenotypes through robust immunophenotyping at a single site. This highly standardized protocol was applied to samples from patients with autoimmune/inflammatory diseases. It is currently used for characterization of the impact of age and environmental factors on peripheral blood immune phenotypes of >400 donors from the initial MI cohort.

## INTRODUCTION

The human immune system provides diverse defense mechanisms against infectious pathogens and tumors. To study the complexity of such responses, numerous systems immunology projects have been developed in the last decade to perform in-depth characterization of immune subpopulations in large cohorts of healthy subjects. These studies aim to identify hereditary and non-hereditary determinants of variation in immune responses (1) and provide resources for identifying biomarkers of disease. Some examples include the Human Immunology Project Consortium (HIPC)(2); the European Network for Translational Immunology Research and Education (ENTIRE)(3); the 10 000 Immunome project (4); the Functional Genomics Project (5); the SardiNIA project (6), and the Milieu Interieur Consortium (MI) (7).

*The Milieu Intérieur* was initiated in 2011 with the aim of establishing the boundaries of a healthy immune response and identifying determinants of immune variability (7). As part of the initial phenotyping of the 1000 healthy donor cohort we developed (8) and applied ten 8-parameter flow cytometry panels to whole blood using a MACSQuant cytometer (Miltenyi Biotec). This analysis allowed us to identify genetic and environmental factors shaping circulating immune cell populations (9). To compare individual versus population aging effects on immunity we recently initiated a 10-year recall study of the original cohort. For cellular phenotyping of this new cohort, we took advantage of recent developments in multi-parameter flow cytometry.

These developments include high-dimensional conventional cytometers (e.g. Symphony A5, BD), new-generation mass cytometers (CyTOF), and recently developed spectral flow cytometers (Aurora, Cytec and ID7000, Sony Biotechnology)(10,11). These advances have been accompanied by the development of new fluorescent dyes and metal-tagged antibodies, which together led to an unprecedented dimensionality in the analysis of cell phenotypes. Spectral flow cytometry offers certain advantages as it allows for 1) simultaneous use of fluorophores of closely emitting spectra, incompatible with conventional flow cytometry; 2) expansion of immunophenotyping panel complexity to 40 parameters and beyond; and 3) measurement of cellular autofluorescence as a separate parameter to avoid false-positive signals. In this study, we describe the development of two 37/36-parameter panels for spectral cytometry that allow standardized immunophenotyping of major immune cell populations in human peripheral blood. In comparison to conventional methods, our approach is less time-consuming and allows for an in-depth analysis from low sample volumes, identification of rare cell subsets and weakly expressed antigens. It is highly compatible for applications to large population-based clinical and translational studies.

## RESULTS

### Choice of cytometer and Panel design

We selected the ID7000™ Spectral Cell Analyzer (Sony Biotechnology) based on a variety of technical features. The instrument has a unique sensitivity in detecting dim signals and rare cell populations and has an integrated deep-well plate reader. In addition, active mixing and cooling of the sample during the acquisition ensure stable acquisition flow rate and preservation of tandem dyes. Our ID7000 is configured with 6 lasers (320 nm, 355 nm, 405 nm, 488 nm, 561 nm, and 637 nm) and 184 detectors. Thanks to its capacity to efficiently identify and separate the dyes with near peak emission signals, spectral cytometry allowed us to characterize all immune cell phenotypes identified in the original MI study (9), and to add several other cell populations such as hematopoietic stem cells (HSC) in just two spectral panels.

We designed two complementary 35 and 34 fluorescent marker cytometry panels to enable identification of major innate and adaptive immune cell populations in 200μl of fresh blood each. Cell proportions, counts, the level of their surface antigen expression and their activation state were measured. A dead cell marker was included in both panels to permit gating on live cells. During the establishment of the panels, 106 different antibodies from four suppliers (BD Bioscience, Sony Biotechnology, ThermoFisher Scientific, Miltenyi Biotec) were tested, of which 43 were excluded (Supplementary table 1). All antibodies were titrated (in 3-5 dilutions) to fit the experimental conditions described in the protocol. Our selection criteria for panel validation were (a) specificity of the signal; (b) signal resolution; (c) availability of the desired fluorochrome; (d) fluorochrome stability (tandem dyes); and (e) price and availability of a single lot of reagents for a cohort study. The specificity of signal was evaluated based on the staining index, i.e., the difference between the positive and the negative populations and the spread of the negative population. Due to the high dimensionality of the panels, we were obliged to include three custom-made antibodies in the innate panel, to match the available fluorescent channels. To minimize the variation of fluorescent signal intensity, only one batch of each antibody was used for staining throughout the whole study. The innate and adaptive panels developed in the study are depicted in Figure 1A.

**Figure 1.**
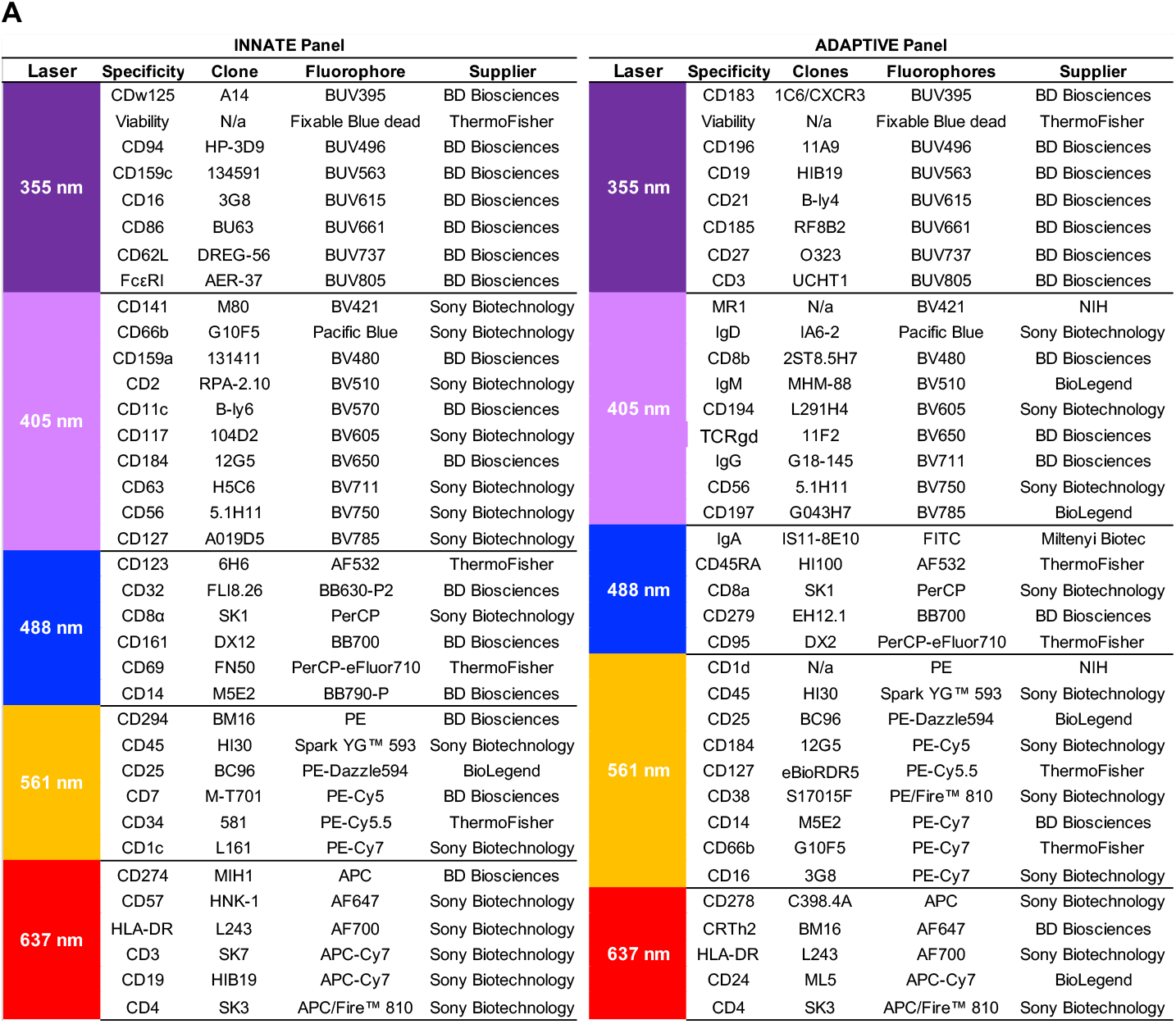
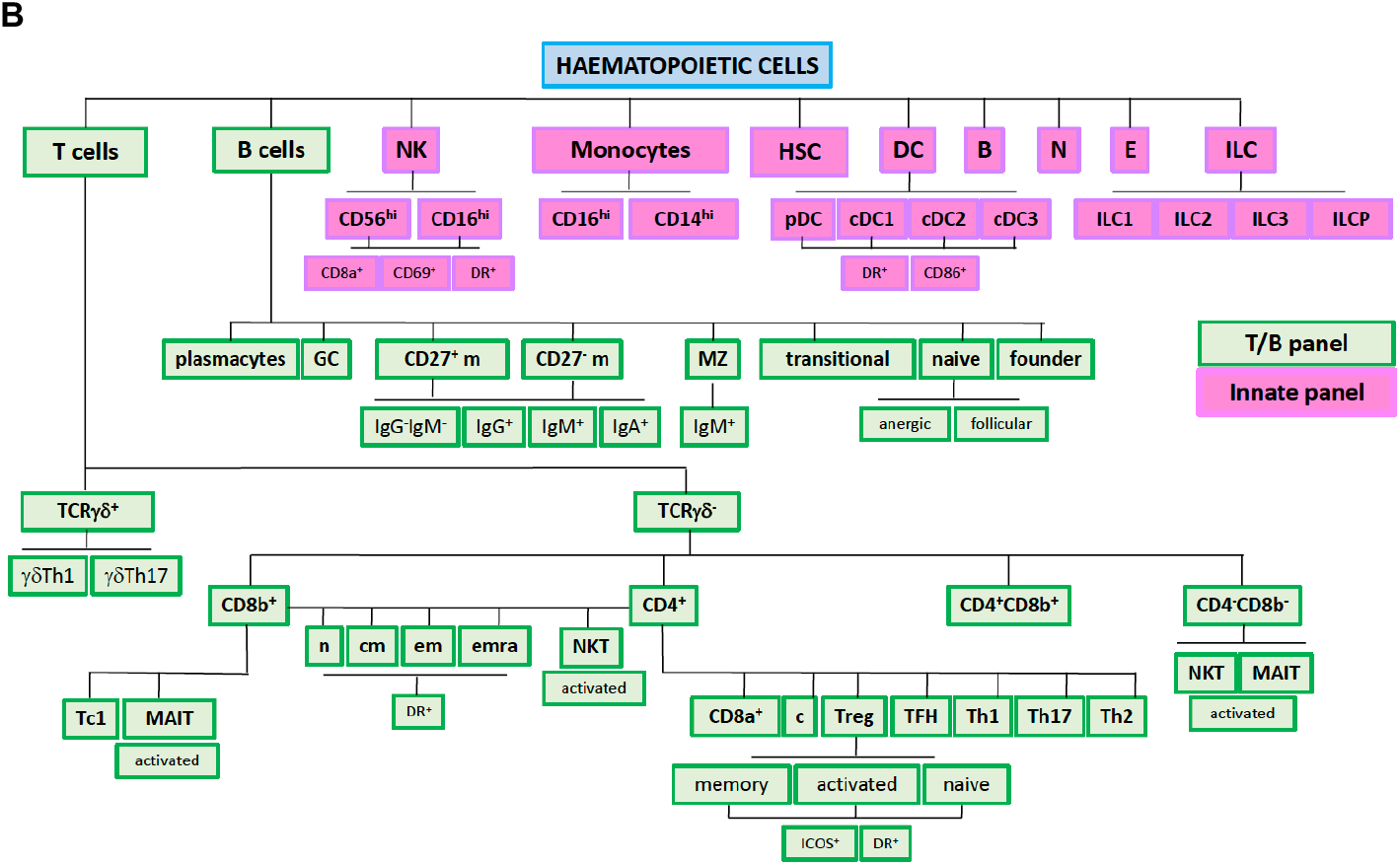
Innate and adaptive panels for identification of major innate and adaptive immune cell populations in fresh peripheral blood. a) 35- and 34-fluorescence panels were designed to enable characterization of cell phenotypes in 200 μl of fresh blood each. b) The scheme represents pre-defined immune cell sub-populations that can be identified. Staining protocol and choice of antibodies enable the identification of 182 immune cell phenotypes.

### Operating procedures

#### Sample preparation and acquisition

The staining protocol was modified from the initial MI study for immunophenotyping of 200μl of fresh whole blood and the acquisition step was standardized, as detailed in the Material and Methods section. We have not observed an added value of blocking Fc receptors (data not shown) and have thus not included this step in our protocols. Since the ID7000 spectral cytometer does not count cells, counting beads (123count eBeads™ Counting Beads, Invitrogen) were used to obtain absolute cell numbers. To each well, 50μl of bead suspension was added prior to the manual mixing step ahead of acquisition.

### Data analysis

#### Gating strategies

In spectral analysis, signals from all detected channels are used to create one spectral emission signal, regardless of the number of fluorochromes analyzed. The unmixed data is typically visualized as a series of images or spectra, each representing the contribution of a single dye to the overall signal. Therefore, the first step of data analysis consists of an unmixing procedure based on spectral libraries that enables identification of each individual fluorophore from a complex spectral signal of multiplexed dyes. The ID7000 uses the WLSM (Weighted Least Squares Method) fluorescence unmixing algorithm to separate the individual spectral fingerprints. Unmixed data are then converted to an FCS-compatible format. The gating strategy for both panels was validated using ID7000 and FlowJo software (version 10.9). We created FlowJo data analysis templates to ensure standardized gating strategy for both panels. The same software was used for manual data analysis that allowed characterization of over 182 cellular subsets and cell activation states. The scheme of all immune cell subpopulations and their phenotypes that can be identified with the two panels is presented in Figure 1B.

#### Innate panel

For characterizing major innate immune cell populations, we first identified CD45^+^ hematopoietic cells, and then excluded doublets using FCS-A/FSC-H and SSC-A/SSC-H (Supplementary figure 1A). Subsequently, T and B cells (CD3^+^ and CD19^+^) were excluded from the subset of live cells to focus on innate subsets. CD16 and CD66b markers allowed us to identify eosinophils and neutrophils (Supplementary figure 1B). Eosinophils were gated within CD66b^+^CD16^-/low^ population as CD123^+^CDw125^+^ cells, and neutrophils were gated as CD66b^+^CD16^+^ cells (Supplementary figure 1C). The activation status of 3 subsets of granulocytes (gating strategy for basophils is described below) was assessed by their expression of CD16, CD32, CD63, CXCR4, FcεRI, HLA-DR, CD62L and PD-L1 molecules.

The CD16^-^ cell population was further distinguished as CD7^+^ or CD7^-^. CD7^+^ population was subsequently identified as NK cells if CD16^+^CD56^+^ and divided into two distinguished subsets (CD56^dim^CD16^bright^, CD56^bright^CD16^dim^) that were additionally characterized by their activation/inhibition status through the expression of CD56, CD69 and CD8a molecules (Supplementary figure 1D). To identify ILCs, the CD16^-^CD56^-^ population was analyzed for the expression of NKG2A and CD94, and the double negative population further for CD127 expression. ILCs were identified among CD127^+^ cells as CD161^-/low^CD25^+^ and subdivided, by expression of CD117 and CRTh2, into ILC2 (CRTh2^+^CD117^-/low^), CD117^-^CRTh2^-^ and CD117^+^CRTh2^-^ (Supplementary figure 1D). Further gating of the CD7^-^ population identified monocytes as CD14^+/-^CD16^+^ that were additionally subdivided into classical (CD14^+^CD16^-/low^), non-classical (CD14^dim^CD16^+^), and intermediate (CD14^med/+^CD16^+^) monocytes. We also assessed the expression of HLA-DR, CD4 and PD-L1 molecules within these populations as markers of activation (Supplementary figure 1E).

In the population of CD14^-^CD16^−^ cells, HLA-DR^+^CD14^-^ cells were selected to subsequently segregate pDCs (CD123^+^CD11c^-^) from cDCs (CD11c^+^CD123^-^). cDCs were additionally divided into CD141^+^ cDCs and CD1c^+^ cDCs as CD141^+^CD1c^-^ and CD141^-^CD1c^+^, respectively (Supplementary figure 1F). The activation status of the three DC subsets was assessed by their expression of HLA-DR, CD4, CD8a, CXCR4 and the costimulatory molecules CD86 and PD-L1.

The CD14^-^ HLA-DR^-^ population resolved by CD45, and CD123 staining was used to identify basophils as CD45^lo^CD123^+^ cells that were further gated as CD123^+^FcεRI^+^ cells. Stem cells were identified as not CD45^lo^CD123^+^ and then as CD34^+^CD117^+^ (Supplementary figure 1B).

#### Adaptive panel

Upon gating on CD45^+^ hematopoietic cells and the exclusion of doublets, the innate compartment (CD14, CD16 and CD66b) was excluded from the viable cell population. CD3 / CD19 gating allowed the discrimination of B (CD19^+^) and T (CD3^+^) cells (Supplementary figure 2A).

T cells were further segregated into TCRγδ positive and negative cell populations. TCR γδ-were characterized as MAIT cells if positive for MR1 tetramer staining and subsequently defined as CD8^+^ or CD4^+^ depending on CD8β and CD4 expression. The non MAIT cells were gated on CD1d to identify NKT cells (CD3^+^CD1d^+^) that were further divided into CD8^+^ and CD4^+^ NKT cells based on the expression of CD8^+^ or CD4^+^ molecules, respectively (Supplementary figure 2B). The activation status of both NKT and MAIT cells were determined by the expression of HLA-DR molecule.

CD4^+^ and CD8b^+^ T cells were gated and analyzed upon exclusion of NKT cells. We characterized naïve (T_N_), central memory (T_CM_), effector memory (T_EM_) and effector memory expressing RA (T_EMRA_) subpopulations of both T cell subsets, based on their expression of CD45RA and CD27 (Supplementary Fig.2B). Since T_N_ and T_CM_ cells have also been defined by the expression of CCR7, we assessed the expression of CCR7 by these cell populations. The activation status was additionally established by the expression of HLA-DR, PD1 and CD56 molecules.

Regulatory T cells (Treg) were identified among CD4^+^ T cells as CD25^+^CD127^-^ and subsequently divided into naïve, memory and activated based on CD45RA and HLA-DR expression (CD45RA^+^HLA-DR^-^, CD45RA^-^HLA-DR^-^, CD45RA^-^HLA-DR^+^, respectively) (Supplementary figure 2B). All 3 Treg subpopulations were additionally investigated for expression of the costimulatory molecule ICOS.

CD25^-^ CD4^+^ T cells were additionally gated on CD127 and four subpopulations (T_N_, T_CM_, T_EM_ and T_EMRA_) were identified among CD127+ cells by expression of CD45RA and CD27 (the same gating strategy as for the CD8^+^ T cells). Expression of CCR7, HLA-DR, PD1, CD56 and CD95 (for T_N_) was subsequently determined. In addition, the CD4^+^ T cell population expressing CD8a was identified (Supplementary figure 2B). The CD127^+^ CD4^+^ T cells expressing the chemoattractant receptor-homologous molecule and/or different chemokine C receptors were identified by two distinct gating strategies: 1) CD45RA^-^CXCR5^+^ were gated on CCR6 and CXCR3 and CXCR5^+^CCR6^-^ subsequently on CXCR3 and CCR4; 2) CD45RA^-^CXCR5^-^ were gated on CCR6 and CXCR5 and CCR6^-^ cells additionally on CXCR3 and CCR4 with subsequent gating of CCR4^+^CXCR3^-^ cells on CRTh2 and CXCR3.

CD19^+^ B cells were gated on CD27 and IgD to detect CD27^+^IgD^+^, CD27^+^IgD^-^, CD27^-^IgD^+^ and CD27^-^IgD^-^ subpopulations. All four subtypes were analyzed for the expression of IgG, IgM and IgA (Supplementary figure 2C). The CD27^+^IgD^-^ memory B cell subtype was furthermore characterized in more detail. Additional gating on CD38 and CD24 allowed segregation of plasmacytes (CD38^high^), germinal center B cells (CD38^low^) and marginal zone B cells (CD24^+/-^ CD38^-^). Marginal zone B cells were further separated into CD24^high^ (CD21^+/low^), CD24^int^ (CD21^+/low^) and CD24^low^ (CD21^+/low^) (Supplementary figure 2C). CD27^-^IgD^+^ B cells were additionally gated on CD38 and CD24 to discriminate naïve B cells (CD38^-^), transitional B cells (CD24^+^CD38^+^), and founder B cells (CD38^-^). CD19^+^ B cells were furthermore gated for the expression of the chemokine receptor CXCR4.

### Assay validation: technical replicates and robustness of the staining procedures

#### Repeatability

As this protocol has been designed for the study of immunological variance in a large cohort (MI 10-year follow-up study), we needed to define the technical variance in our immunophenotyping procedures by performing reproducibility and repeatability experiments. To confirm the repeatability, we analyzed the same sample, in five independent runs, by a single operator. In our experimental setting, fresh blood samples from five healthy donors were separated into five aliquots each then processed and stained separately using each panel, as described in the operating procedure. The results of this test for both panels were highly consistent, with intra-panel coefficients of variation (CVs) below 15% for most of the analyzed cell subsets, irrespective of their absolute counts (Figure 2). For the major circulating immune populations, including neutrophils, eosinophils, basophils, monocytes, T cell subsets (Treg, CD4^+^, CD8^+^, MAIT), and B cells, CV values were between 1.09% and 9.13% (cell proportions) and between 4.33% and 14.82% (absolute cell numbers). This was the case even for the rare cell populations like cDCs (14.51% and 7.25%, cell proportion and absolute cell number, respectively), pDCs (9.52% and 13.72%, cell proportion and absolute cell number, respectively) and ILCs (11.75% and 11.68%, cell proportion and absolute cell number, respectively).

**Figure 2.**
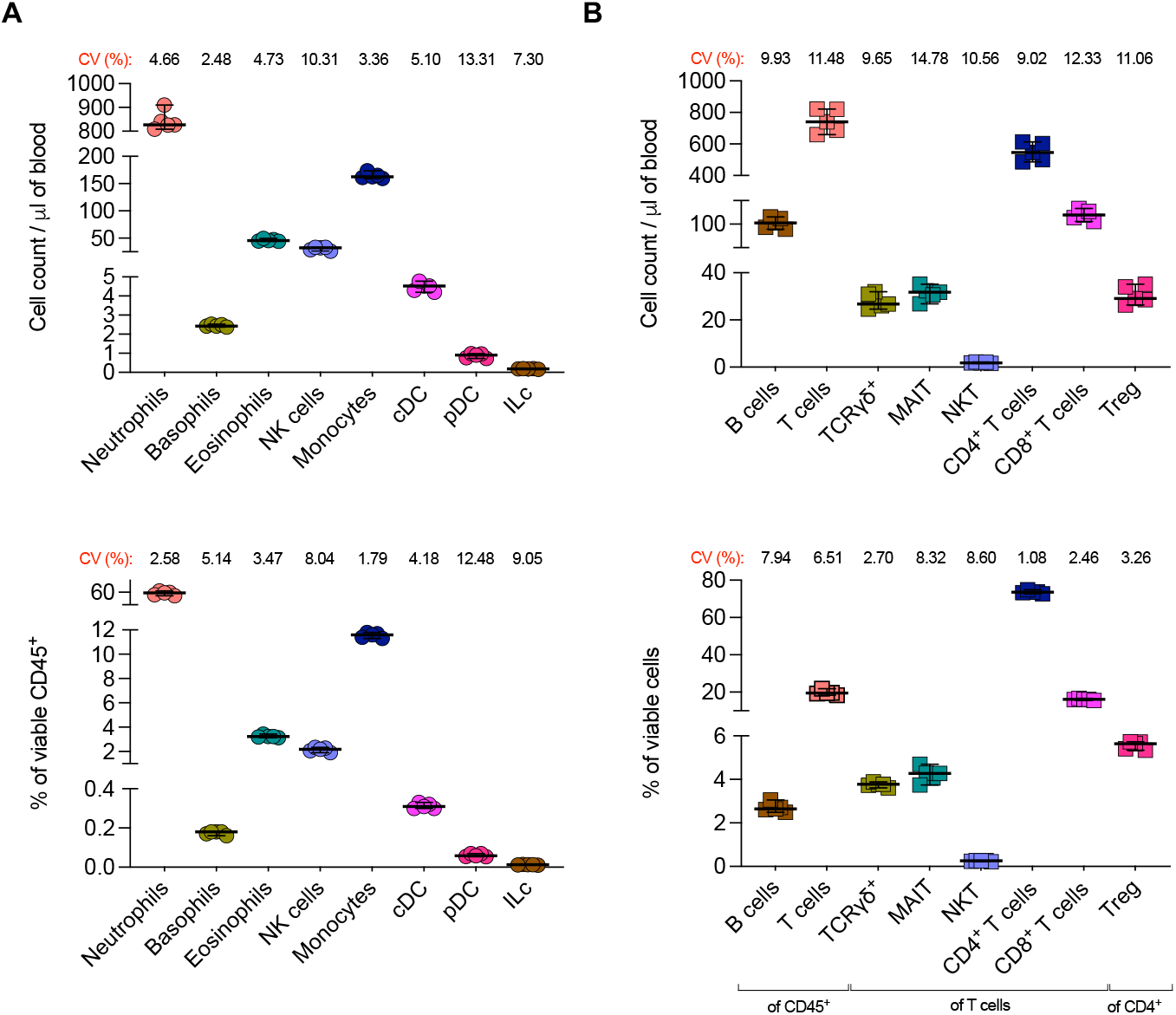
Repeatability study using fresh whole blood. Fresh blood samples of a donor were aliquoted in five tubes with 200 μl of blood, processed and stained by innate and adaptive panels, as described. This was done for five donors in total. Numbers (upper panels) and proportions (lower panels) of selected innate (A) and adaptive (B) cell subsets are shown. The CVs for serial measurements are indicated for each analyzed immune cell population.

#### Reproducibility

To confirm the robustness of the staining procedures and the stability of staining over time, an important consideration for large cohort studies, we assessed reproducibility of our assays. To provide a stable reference, we utilized commercially available stabilized human blood (CD-Chex Plus BD, Eurobio Scientific) that was analyzed in five independent experiments, across a 2-week period (Figure 3). These data showed reproducible results with CVs under 13% for proportions of NK cells (4.37%), monocytes (7.64%), ILCs (12.44%) and neutrophils (3.55%), and CVs under 15% for proportions of B cells (14.30%) and T cell subsets (TCRγ δ (9.03%), CD4^+^ (3.64%), CD8^+^ (3.93%)).

**Figure 3.**
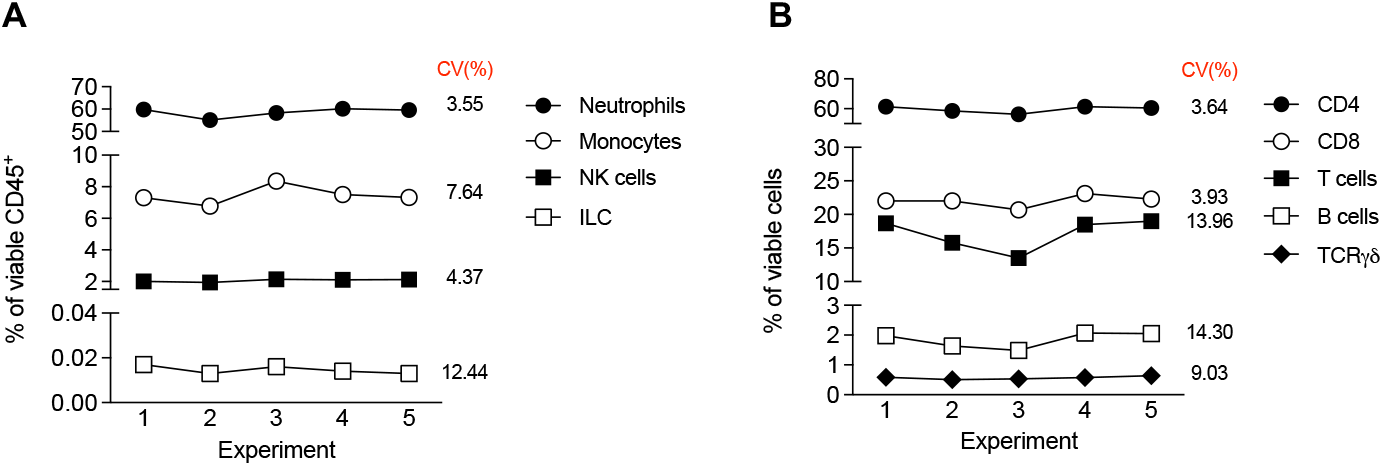
Reproducibility assay using commercially available stabilized blood. Stabilized human blood (CD-Chex Plus, Eurobio Scientific) was analyzed in five independent experiments, across a 2-week period. The percentages of indicated cell subsets for (A) innate and (B) adaptive panels are shown. The CVs for serial measurements are indicated for each analyzed immune cell population.

#### Fresh blood vs PBMCs

For certain studies and clinical trials, accessibility of fresh whole blood for flow cytometry analysis can be logistically challenging. The most common solution is density gradient centrifugation and isolation of peripheral blood mononuclear cells (PBMCs) and subsequent freezing, which can introduce technical variability (12) and remove important immune cell subsets (e.g. granulocytes). More recent technical solutions (e.g. Cytodelics, SMART tube) allow freezing of whole blood for later analysis (13). Given our interest to phenotype granulocytes, removed during PBMC isolation, we tested the Cytodelics blood stabilization kit. Unfortunately, the fixation led to loss of signal for certain antibodies, likely due to conformational changes of targeted epitopes (Supplementary table 2). Given the necessity of some studies to work with PBMCs, we also tested our panels on freshly isolated PBMCs and PBMCs preserved in CellCover solution and compared the results with those of fresh whole blood from the same donors, obtained in the same experimental setup. The comparison was made between 200μl of fresh blood, 1 million PBMCs and 0.2 million PBMCs. While 0.2 million PBMCs correspond to the estimated number of PBMCs in 200μl of fresh blood, a sample with 1 million PBMCs was included to allow the detection of minor and rare cell populations or phenotypes with higher precision. This also allowed us to test our staining conditions with 5-fold superior cell numbers compared to what it was originally developed for, and to confirm that our panels and staining procedures work equally well with a broad range of samples.

As can be seen in Figure 4, the percentage of viable cells and absolute cell numbers between whole blood and corresponding PBMCs were comparable for both panels. As expected, the percentage of viable B and T cells was higher in PBMCs than in corresponding whole blood (Figure 4A), while the absolute cell numbers were not significantly different (Figure 4B). Certain differences were observed for NKT, Treg and TCRγδ cells, which might be due to low population proportions and/or cell numbers. Regarding the innate panel, as expected, neutrophils and eosinophils were well detected in whole blood and, to a lesser extent in PBMCs (Figure 4C-D), whereas the percentage of viable basophils was higher in PBMCs than in whole blood (Figure 4C). This was not the case when absolute numbers were compared (Figure 4D). The proportion of NK cells, monocytes, cDCs, pDCs, and ILCs among the viable cells were at comparable levels (Figure 4C), while absolute numbers for NK cells, cDCs, and pDCs were significantly higher in fresh blood (Figure 4D). This could be explained by cell loss during the PBMC isolation procedure, which may be higher for rare cell populations.

**Figure 4.**
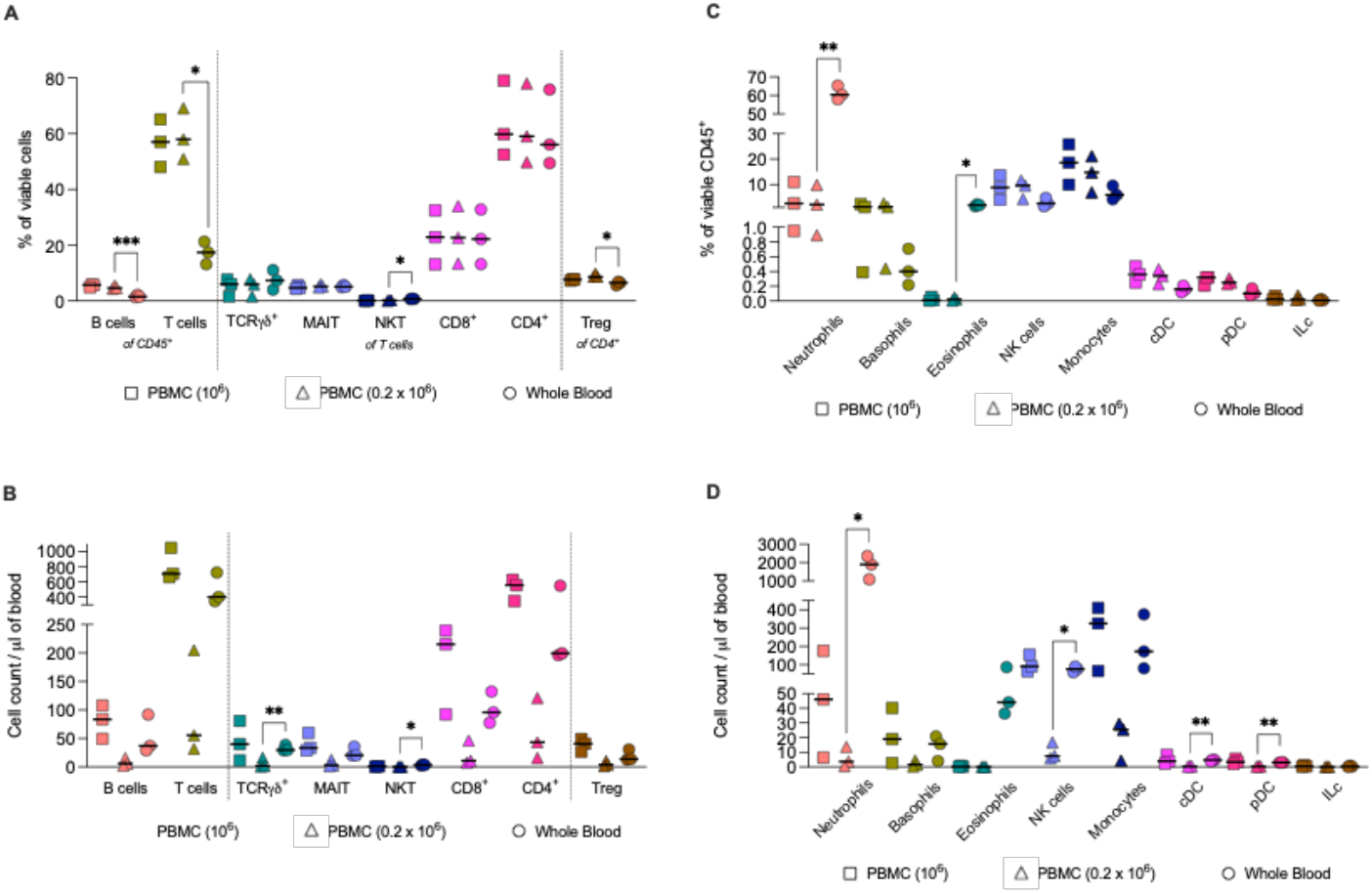
Fresh blood vs PBMC comparison. Fresh blood from 3 donors was partly used for PBMC isolation and both types of samples were analyzed in a single experiment. The percentages of indicated cell subsets for (A) adaptive and (C) innate panels and absolute cell numbers for (B) adaptive and (D) innate panels are shown. The significant difference between 0.2 million PBMCs and fresh blood was expressed by the *. *P* values were calculated by paired t-test. *, P<0.05; **, P<0.005; ***, P<0.0005.

### Semi-automation

The implementation of automated procedures can help to eliminate possible errors or variation caused by repetitive work during a prolonged period by technical personnel, which is particularly relevant for clinical studies with large sample numbers. Therefore, we took advantage of automation in sample preparation for our standardized cellular immunophenotyping protocol. To achieve this, we implemented our protocol using the Freedom EVO150 liquid handling platform (Tecan). The premix of antibodies was prepared manually on a daily basis, and all other steps for the sample preparation and staining protocol were performed using the liquid handling platform, EVO150, with the exception of centrifugation. The pipetting scripts for the platform were created to enable the staining of 10 samples, in parallel, in 5 ml round bottom tubes. The script is available at https://github.com/cbutechs/MI_spectral. The automation process requires validation prior to implementation in clinical studies. To this end, one operator performed a repeatability assay within a single day manually and with the automation platform in parallel. One whole-blood sample from a healthy donor was divided into ten aliquots and immediately processed and stained in parallel (5 manually, 5 automated). Results from representative analysis of three experiments are shown in Figure 5. For all analyzed immune cell populations, no significant difference between absolute cell numbers was observed, when comparing manual experimentation with the semi-automated platform. This was the case for major cell populations such as B cells (P=0.0625), T cells (P=0.1250), neutrophils (P=0.1875), monocytes (P=0.6250), but also for minor immune cell populations such as pDCs (P=0.6250), ILCs (P=0.6250), MAIT cells (P=0.1250), NKT cells (P=0.0625) and Treg (P>0.9999).

**Figure 5.**
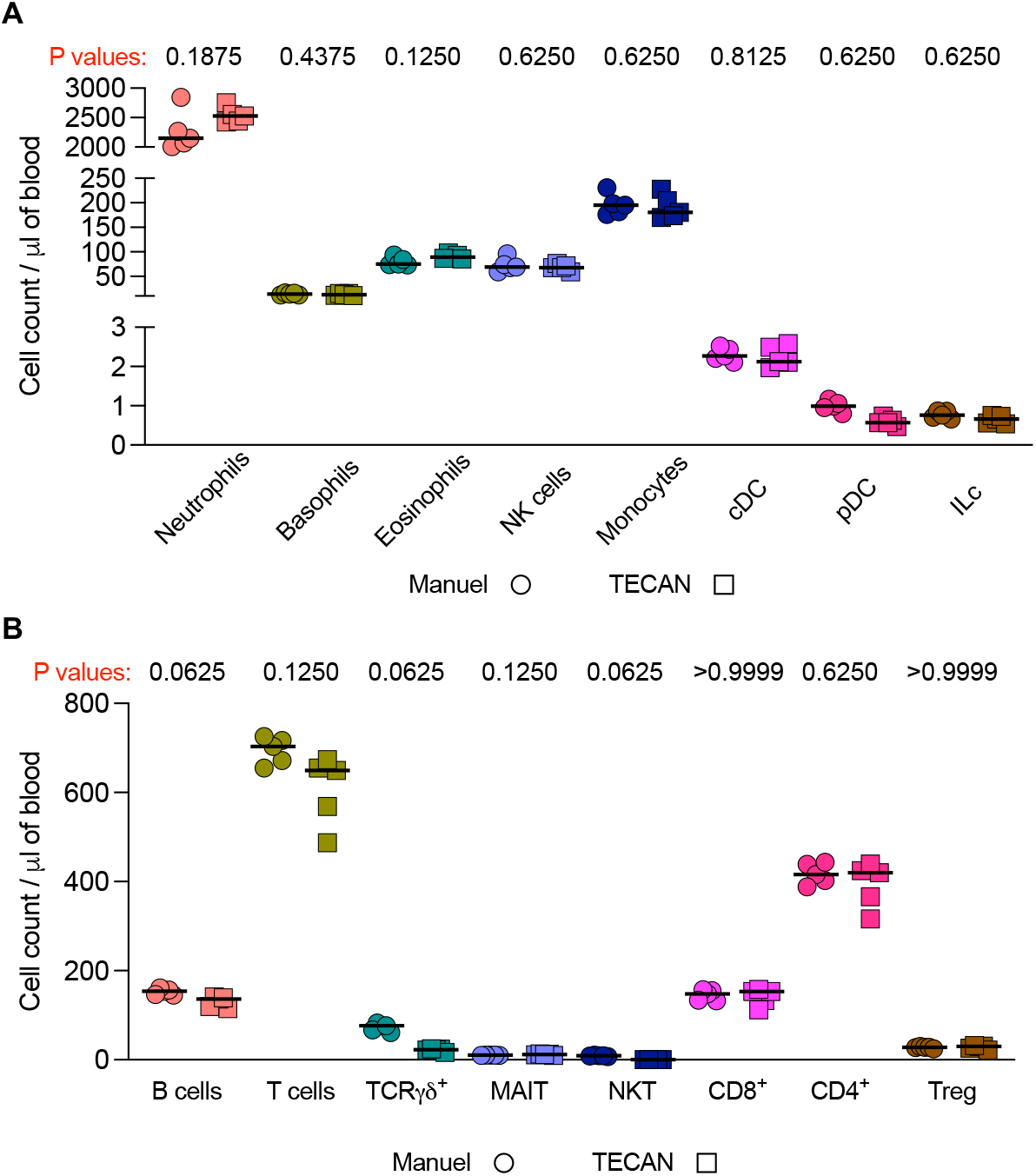
Semi-automation of sample processing and staining of whole blood. Fresh blood samples of a donor were aliquoted in five tubes with 200 μl of blood, processed and stained by innate and adaptive panels, manually and with automated platform in parallel. This was performed for three donors. Numbers of selected innate (A) and adaptive (B) cell subsets are shown. The difference between the two methods was expressed by the P value indicated for each analyzed immune cell population. *P* values calculated by Wilcoxon matched-pairs signed rank test.

### Application to phenotyping of samples from patients with autoimmune/inflammatory diseases

We wanted to explore whether our panels and method are sensitive enough to capture differences in immune cell phenotypes between healthy subjects and subjects with pathological conditions. For this proof-of-concept application, we compared a small number of patients with different autoimmune/autoinflammatory conditions (n=8) (Supplementary table 3) with age and sex matched healthy donors (n=8). From 182 phenotypes compared using the two panels, 16 minor/rare cell populations including NKT, CD1c^+^ cDCs, MAIT and IgG^+^IgM^+^CD27^-^IgD^+^cells showed significant differences with an uncorrected p value of <0.05. Four phenotypes, all related to B cell subsets, remained significant even after correcting for multiple testing (Figure 6). We have not further explored the observed differences since it was out of the scope of this work. Nevertheless, this validation step confirmed the advantage and suitability of our protocol for clinical trials or studies that require the detection of minor changes of rare immune cell populations and their phenotypes introduced by different pathological environments.

**Figure 6.**
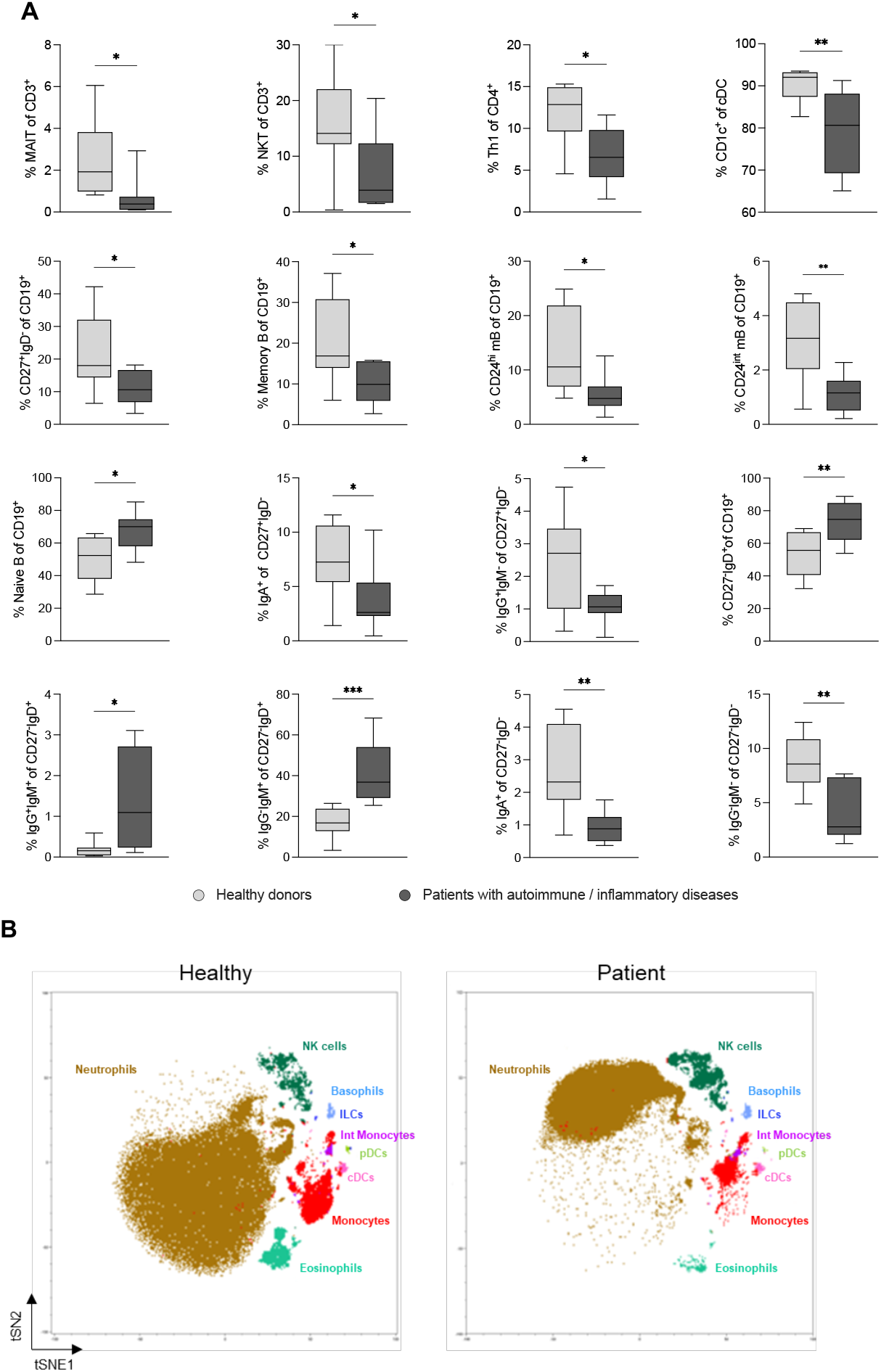
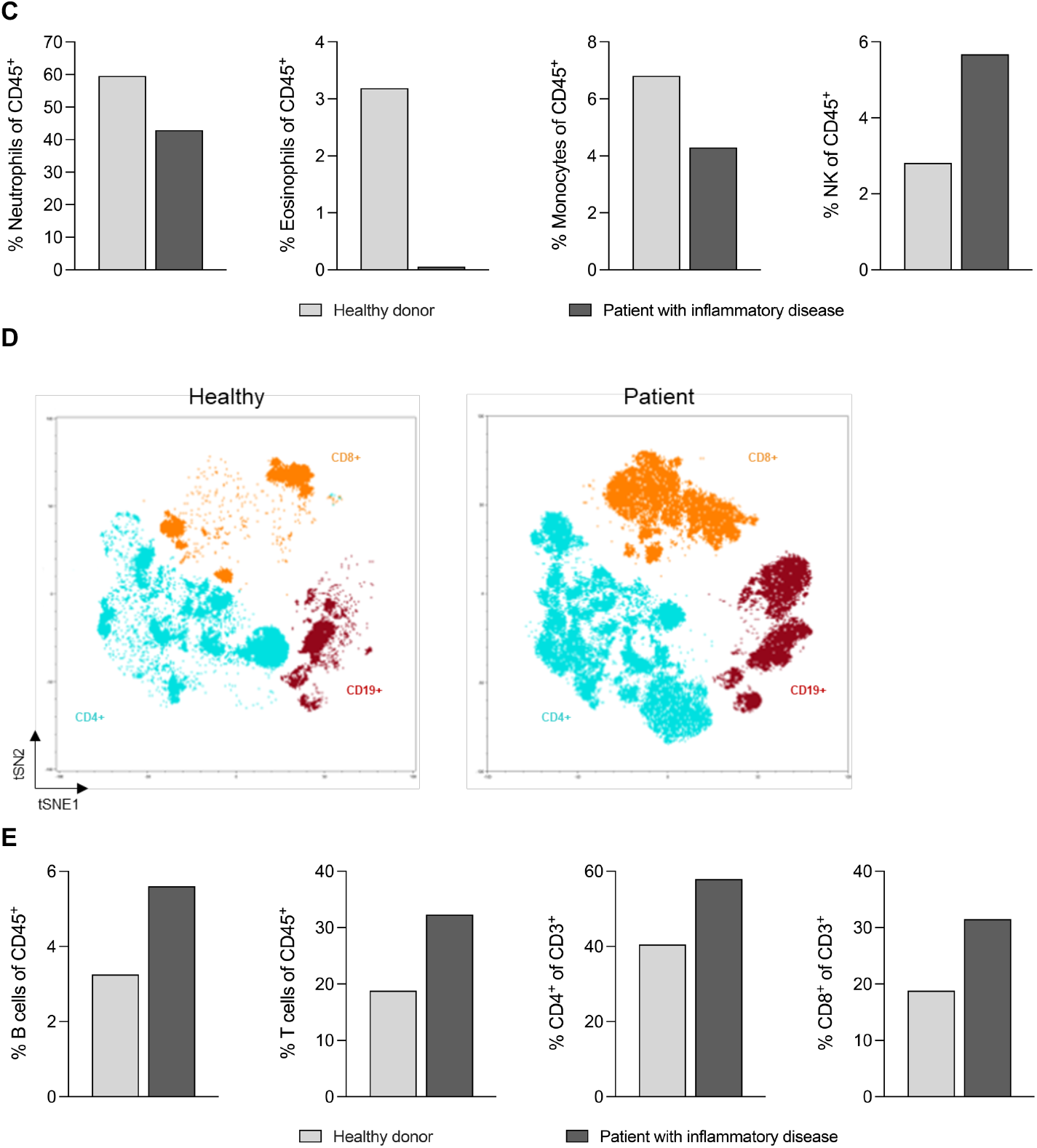
Panel validation in patient samples. Fresh blood from 8 patients with autoimmune/inflammatory disease was analyzed and compared with age/sex-matched donors from MI 10-year follow-up study. (A) Phenotypes characterized by supervised (FlowJo) analysis. The unpaired t-test was used for statistical analysis: *, P<0.05; **, P<0.005; ***, P<0.0005. (B, D) An unsupervised analysis was performed by the prototype of Sony software. The input data were single alive dump negative cells. Data from one healthy and one patient subject were concatenated prior to the unsupervised analysis. We used 750 iterations and the perplexity of 30 as parameters for creating the Flt-SNE representations. KNN algorithm was used for clustering. Shown are tSNE projections of clusters annotated based on manual gating. A representative example of a healthy donor (left panels) and of a patient with an inflammatory disease (right panels), is shown for (B) Innate panel and (D) adaptive panel. (C, E) Supervised analysis (FlowJo) of the same subjects and the same immune cell population as for non-supervised analysis, for (C) Innate panel and (E) adaptive panel.

### Unsupervised data analysis

Analysis of high-dimensional flow cytometry data is the major challenge of advanced cell phenotyping. We assessed the added value of data visualization after non-supervised analysis. To this end, in addition to FlowJo analyses, we employed a non-supervised approach to visualize major cell subsets identified by innate and adaptive panels in one patient with inflammatory disease and one age/sex-matched healthy donor. We used a prototype of Sony data analysis software that relies on algorithms like FIt-sne, UMAP and flowAI incorporated in the native prototype Sony Software. It uses the original Sony ID7000 format files instead of FCS.

As shown in Figure 6B, there were obvious differences between major innate immune cell populations (e.g., neutrophils, eosinophils, monocytes, NK cells). Interestingly, the Flt-sne representation indicated a distinct neutrophil distribution between healthy and patient subjects, suggesting phenotypic differences within this cell population that could be potentially further explored (not within the scope of this study). An example of non-supervised analysis for adaptive cell subsets is shown in Supplementary figure 3. Evident differences between a patient with inflammatory disease and a sex/age-matched healthy donor can be seen for CD4^+^ T cells (naïve, CM, EM, EMRA) (Supplementary figure 3A), CD4^+^ T cell subsets (Th, Treg, MAIT) (Supplementary figure 3B), CD8^+^ T cells (naïve, CM, EM, EMRA) (Supplementary figure 3C) and B cell subsets (Supplementary figure 3D).

To confirm that discrepancies between the two subjects were not a result of technical bias introduced by the non-supervised analysis tool, we compared these results to manual analysis of these two samples. As shown in Figure 6C for innate panel, the results of manual analysis matched those obtained by non-supervised approach. Similar was obtained for adaptive panel (B cells, T cells, CD4^+^ T cells, CD8^+^ T cells) as shown for non-supervise (Figure 6D), compared to supervised analysis (Figure 6E).

## DISCUSSION

Multi-parameter flow cytometry is a powerful technique that allows deep characterization of immune cell subpopulations in clinical and translational studies. However, a major challenge is the correct choice of tools and techniques that allow reliable and performant comparison between individual subjects and across different studies. To this end, the standardization and development of robust protocols is of the highest importance.

In 2011, we developed a standardized procedure for flow cytometry allowing us to analyze the 1,000 healthy donors of the *Milieu Intérieur* study. To analyze major immune phenotypes and considering the technological advances at that time, we established a standardized staining protocol with 10 separate panels. Over the past years, flow cytometry technologies have significantly improved, resulting in the development of new, more powerful cytometers such as spectral cytometry, first commercialized by Sony Biotechnology (10). The simultaneous development of new reagents, monoclonal antibodies, and a variety of new fluorochromes allowed us to replace ten 8-color panels with two 35-34-color panels. Consequently, the benefit is multiple: a) decrease in required sample volume, b) considerable time reduction and therefore c) cost reduction.

Spectral flow cytometry is heavily used in the field of immunology, where there is a need for simultaneous analysis of as many cell markers as possible to characterize different immune cell populations and their phenotypes. We considered the CytoFLEX (Beckman Coulter) and Symphony A5 (BD Bioscience) flow cytometers for this project and performed the initial testing of panels on these machines. However, the obtained results were not satisfactory. Although equipped with a deep-well plate reader, CytoFLEX was limited by low dimensionality. The high dimensionality of our panels rendered the complexity of panel design that was challenging for the Symphony A5. In addition, the instrument lacked the possibility to perform the acquisition from Deepwell plates.

One limitation of the ID7000 is its lack of automated cell counting. While in the initial MI study we relied on a cytometer that had a cell counting capacity, here, we were dependent on the use of counting beads to assess absolute cell numbers. This resulted in less satisfying CVs for the reproducibility and repeatability tests, in particular for minor/rare immune cell subsets.

The major challenge of high-dimensional flow cytometry (conventional and spectral) remains data analysis. It starts with compensations in the case of conventional cytometry, and with spectral signal unmixing in spectral cytometry. Both have a significant impact on the results and reproducibility. The manual unmixing and its adjustments is both time-consuming and biased by its manual nature. The proprietary data format for the raw data makes finding automated procedures challenging. Traditional analysis using 1 and 2 dimensional gates is usually done manually and is difficult to reproduce (14). While the gold standard FlowJo approach is convenient for defining an initial gating strategy, application of an unsupervised data analysis pipeline is better suited for the large number of measured parameters (>35) and for the analysis of numerous samples in clinical studies (15). The unsupervised approach allows identification of novel cellular clusters, without prior knowledge of their characteristics, potentially leading to the discovery of previously uncharacterized immune cell populations and their roles in various disease states. Most methods and workflows currently rely on self-organizing maps (KOHONEN)(16) like FlowSOM (17,18) or Catalyst (19). Several pipelines for unsupervised analysis are commercially available (e.g. OMIQ, Tercen, Ozette, METAFORA, FCS express, Cytobank, FlowJo plugins), but all show certain limitations. Unsupervised, similarly to supervised pipelines imply the choice of transformation and optimization of its parameters (biexponential versus inverse hyperbolic sine (asinh)), for which there is still no consensus in the community. When dealing with human samples, due to the large variability of data, spectra need to be aligned, which complicates the analysis of fluorescent shift. The key is to ensure that the applied method is reproducible and thus adapted to further studies.

The major added value of a standardized protocol like ours is its versatility. We have demonstrated its applicability in the analysis of both fresh blood and PBMCs, which renders it beneficial for a vast range of studies and clinical trials. In addition, we have demonstrated the capacity of our approach to identify small shifts in cell phenotypes in pathological conditions, by applying it to the analysis of samples from a small cohort of patients with autoimmune/inflammation diseases and comparing them to sex/age-matched healthy donors from MI 10-year follow-up study. The obtained results confirm the capacity of our protocol to detect changes (even) in rare, minor cell populations.

In summary, our standardized protocol and two high-dimensional spectral cytometry panels are well-adapted to immunomonitoring in a wide range of clinical studies based on analysis of fresh blood and/or PBMC.

## MATERIALS and METHODS

### Healthy donors

Fresh whole blood was collected from healthy French volunteers enrolled at the Clinical Investigation and Access to BioResources (ICAReB) platform (Center for Translational Research, Institut Pasteur, Paris, France). These donors were part of the CoSImmGEn cohort (NCT03925272). Additional blood was obtained from donors recruited as part of the Milieu Interieur 10 year longitudinal V3 study (NCT05381857) performed at Biotrial, Rennes. This clinical study was approved by the CPP (Comités de Protection des Personnes) Nord Ouest III and ANSM (Agence nationale de sécurité du médicament et des produits de santé). Written informed consent was obtained from all study participants. Stabilized whole blood was obtained from Streck (350174/39).

### Patient samples

Human patient samples were collected in Cochin Hospital (Paris, France), in the setting of the local RADIPEM biological samples collection derived from samples collected in routine care. Biological collection and informed consent were approved by the Direction de la Recherche Clinique et Innovation (DRCI) and the French Ministry of Research (N°2019-3677).

### Sample preparation

Whole blood was collected on Li-heparin as anti-coagulant and maintained at 18–25°C until processing (6 to 8 h). To eliminate soluble antibodies and other molecules that may interfere with staining, 1 ml of whole blood was washed by mixing fresh whole blood and PBS at a ratio of 1:1, followed by centrifugation at 500 g for 5 min at 18-22°C (room temperature, RT). The supernatant was discarded, PBS was added up to 1 ml and shortly vortexed.

### Staining protocol

Washed blood (200 μl per staining panel) was added to Live/dead solution to obtain 1:1000 Live/dead final dilution and incubated for 30 min at RT protected from light. Thereafter, 2 ml of PBS were added to the tubes, centrifuged for 5 min at 500 g, and the supernatant was discarded.

Antibody premixes were prepared, shortly vortexed, spun for 10 s, and added on the surface of the blood pellet. The samples were shortly vortexed and incubated for 20 min at RT, protected from light. 2 ml of PBS were added to the tubes, centrifuged for 5 min at 500 g, and the supernatant was discarded.

All samples were resuspended in 4 ml of RBC lysing solution (BD FACS Lysing Solution, BD Biosciences), vortexed and incubated for 15 min at RT protected from light. Of note, BD FACS Lysing Solution also contains a fixative reagent. Following centrifugation (5 min at 500 g), the supernatant was discarded, the samples were resuspended in 2 ml of PBS to stop the reaction. Upon centrifugation for 5 min at 500 g, the supernatant was discarded, the samples were resuspended in 250 μl PBS, transferred to a 500 μl Deepwell plate and stored at 4°C until acquisition (overnight), protected from light.

### Sample acquisition

Before sample acquisition, the ID7000 cytometer was calibrated using fluorescent calibration beads according to the manufacturer’s instructions. An acquisition template was created for each panel. Spectral cytometry does not require compensation but does require single stained controls to allow unmixing of obtained data. To this purpose we used OneComp eBeads (eBioscience, ThermoFischer) or single stained cells. The latter were used for tetramers and live/dead marker. The unstained sample was used for measurement of the background signal. Prior the acquisition, counting beads (123count eBeads, ThermoFischer) were vortexed for 30 s and 50μl were added to each sample.

To ensure the reproducibility of the protocol, the samples were manually mixed just before the acquisition. In addition, the same order of acquisition of the two panels was executed (innate panel first). For each panel, the acquisition of samples was followed by that of unstained control, and of single-stained controls. To avoid cross-contamination, an additional Priming/Washing step was systematically performed between the acquisition of the two panels. The samples were acquired at a speed of around 100 μl per minute, which represented a good compromise between the number of acquired cells, the acquisition time (3 min), the abort and the saturation rate. The dead volume was between 30 and 50 μl.

A delay of 5 s between the beginning of aspiration and start of sample recording ensured recording of data with stable acquisition rate.

### SData analysis

Spectral cytometry data were generated using ID7000 software version 1.1.12.25251 and saved in exdat format (SONY). The spectral data were unmixed using the WLSM algorithm of ID7000 software. Then, the unmixed sample data were converted to FCS files and analysed using FlowJo version 10.9 software. Statistical graphs were prepared using Graphpad Prism software version 9.5.1.

*The Milieu Intérieur Consortium is composed of the following team leaders: Laurent Abel (Hôpital Necker), Andres Alcover, Hugues Aschard, Philippe Bousso, Nollaig Bourke (Trinity College Dublin), Petter Brodin (Karolinska Institutet), Pierre Bruhns, Nadine Cerf-Bensussan (INSERM UMR 1163 – Institut Imagine), Ana Cumano, Caroline Demangel, Christophe d’Enfert, Ludovic Deriano, Marie-Agnès Dillies, James Di Santo, Gérard Eberl, Jost Enninga, Jacques Fellay (EPFL, Lausanne), Ivo Gomperts-Boneca, Milena Hasan, Magnus Fontes (Institut Roche), Gunilla Karlsson Hedestam (Karolinska Institutet), Serge Hercberg (Université Paris 13), Molly A. Ingersoll (Institut Cochin), Rose Anne Kenny (Trinity College Dublin), Olivier Lantz (Institut Curie), Mickael Ménager (INSERM UMR 1163 – Institut Imagine), Frédérique Michel, Hugo Mouquet, Cliona O’Farrelly (Trinity College Dublin), Etienne Patin, Sandra Pellegrini, Stanislas Pol (Hôpital Côchin), Antonio Rausell (INSERM UMR 1163 – Institut Imagine), Frédéric Rieux-Laucat (INSERM UMR 1163 – Institut Imagine), Lars Rogge, Anavaj Sakuntabhai, Olivier Schwartz, Benno Schwikowski, Spencer Shorte, Frédéric Tangy, Antoine Toubert (Hôpital Saint-Louis), Mathilde Touvier (Université Paris 13), Marie-Noëlle Ungeheuer, Christophe Zimmer, Matthew L. Albert (Hibio), Darragh Duffy, and Lluis Quintana-Murci. Unless otherwise indicated, partners are located at Institut Pasteur, Paris. Darragh Duffy and Lluis Quintana-Murci are co-coordinators of the Milieu Intérieur Consortium.

## Acknowledgements

We thank the ICAReB platform of the Institut Pasteur for healthy donor’s fresh blood samples. We thank BD for providing the antibodies for testing and NIH Tetramer Core Facility for providing the tetramers used in the study. This work benefited from support of the French government’s Invest in the Future Program, managed by the Agence Nationale de la Recherche (ANR, reference 10-LABX-69-01).

