## Supplemental Tables and Figures for "High-dimensional spectral cytometry panels for whole blood immune phenotyping"

| Laser | Specificity | Clone | Fluorophore | Supplier | References | Panel | Notes |
|---|---|---|---|---|---|---|---|
| 355 nm | CD3 | UCTH1 | BUV615 | BD Biosciences | 612993 | Adaptive | 1 |
| | CD38 | HIT2 | BUV661 | BD Biosciences | 612969 | Adaptive | 1 |
| | CD115 | 9-4D2-1E4 | BUV737 | BD Biosciences | 749099 | Innate | 3 |
| | CD203c | NP4D6 | BUV737 | BD Biosciences | 752538 | Innate | 2 |
| | CD27 | M-T271 | BUV737 | BD Biosciences | 741833 | Adaptive | 2 |
| | CD27 | L128 | BUV737 | BD Biosciences | 612830 | Adaptive | 4 |
| | CD21 | B-ly4 | BUV805 | BD Biosciences | 742008 | Adaptive | 1 |
| 405 nm | CD141 | 1A4 | BV421 | BD Biosciences | 565321 | Innate | 3 |
| | CD279 | EH12.2H7 | BV421 | Sony Biotechnology | RT2249595 | Adaptive | 3 |
| | CD24 | ML5 | BV510 | BD Biosciences | 563035 | Adaptive | 1 |
| | CD24 | ML5 | BV510 | Sony Biotechnology | RT2155630 | Adaptive | 1 |
| | IgM | MHM-88 | BV570 | Sony Biotechnology | RT2172585 | Adaptive | 2 |
| | CD194 | 1G1 | BV605 | BD Biosciences | 562906 | Adaptive | 4 |
| | CD117 | 104D2 | BV605 | BD Biosciences | 562687 | Innate | 4 |
| | Integrin β7 | FIB504 | BV650 | BD Biosciences | 564285 | Adaptive | 3 |
| | CD294 | BM16 | BV711 | BD Biosciences | 740813 | Adaptive | 1 |
| | TCR Vα7.2 | 3C10 | BV711 | Sony Biotechnology | RT2358655 | Adaptive | 5 |
| | CD127 | HIL-7R-M21 | BV786 | BD Biosciences | 563324 | Innate | 3 |
| 488 nm | CD45RA | HI100 | BB515 | BD Biosciences | 564552 | Adaptive | 1 |
| | CD336 | p44-8 | BB515 | BD Biosciences | 566025 | Innate | 3 |
| | CD203c | NP4D6 | FITC | Sony Biotechnology | RT2223070 | Innate | 3 |
| | CD79a | HM47 | FITC | Sony Biotechnology | RT2267555 | Adaptive | 3 |
| | CD185 | MU5UBEE | AF532 | ThermoFisher | 58-9185-42 | Adaptive | 3 |
| | TCRγδ | B1 | BB630-P2 | BD Biosciences | **Custom** | Adaptive | 3 |
| | TCRV24 | 6B11 | BB790-P | BD Biosciences | **Custom** | Adaptive | 5 |
| 561 nm | CD294 | BM16 | PE | Sony Biotechnology | RT2350525 | Innate | 3 |
| | CD25 | M-A251 | PE-Dazzle594 | Sony Biotechnology | RT2380630 | Innate & Adaptive | 2 |
| | CD45 | HI30 | PE-AF610 | ThermoFisher | MHCD4522 | Innate & Adaptive | 1 |
| | CD7 | CD7-6B7 | PE-Cy5 | Sony Biotechnology | RT2315550 | Innate | 2 |
| | CD197 | 3D12 | PE-Cy5.5 | ThermoFisher | 35-1979-42 | Adaptive | 3 |
| | CD115 | 9-4D2-1E4 | PE-Cy7 | Sony Biotechnology | RT2336535 | Adaptive | 3 |
| | CD14 | 63D3 | PE-Cy7 | Sony Biotechnology | RT2435555 | Adaptive | 3 |
| | CD16 | 3G8 | PE-Cy7 | BD Biosciences | 560918 | Adaptive | 4 |
| | CD66b | G10F5 | PE-Cy7 | Sony Biotechnology | RT2125575 | Adaptive | 2 |
| 647 nm | CD138 | DL-101 | AF647 | Sony Biotechnology | RT2361565 | Adaptive | 3 |
| | IgG | ICO-97 | AF647 | Novus Biologicals | NBP2-34647AF647 | Adaptive | 3 |
| | CD4 | S3.5 | APC-Cy5.5 | ThermoFisher | MHCD0419 | Innate & Adaptive | 1 |
| | CD3 | SK7 | APC-Cy7 | BD Biosciences | 561800 | Innate | 4 |
| | CD5 | UCHT2 | APC-Cy7 | BD Biosciences | 563516 | Innate | 2 |
| | CD5 | L17F12 | APC-Cy7 | Sony Biotechnology | RT2420050 | Innate | 3 |
| | IgG | HP6017 | APC-Cy7 | Sony Biotechnology | RT2646565 | Adaptive | 3 |
| | IgM | UCH-B1 | APC-Cy7 | Novus Biologicals | NBP2-78050APCCY7 | Adaptive | 1 |
| | CD38 | HIT2 | APC/Fire810 | BioLegend | 303549 | Adaptive | 1 |

**Supplementary table 1.** Antibodies tested during the development of the two panels. The number indicate as follows: 1- Fluorochrome incompatible with the final panel; 2- Unspecific staining in whole blood; 3- No signal; 4- Antibody performing equally well as the selected antibody; 5- Replaced by more specific tetramer staining.

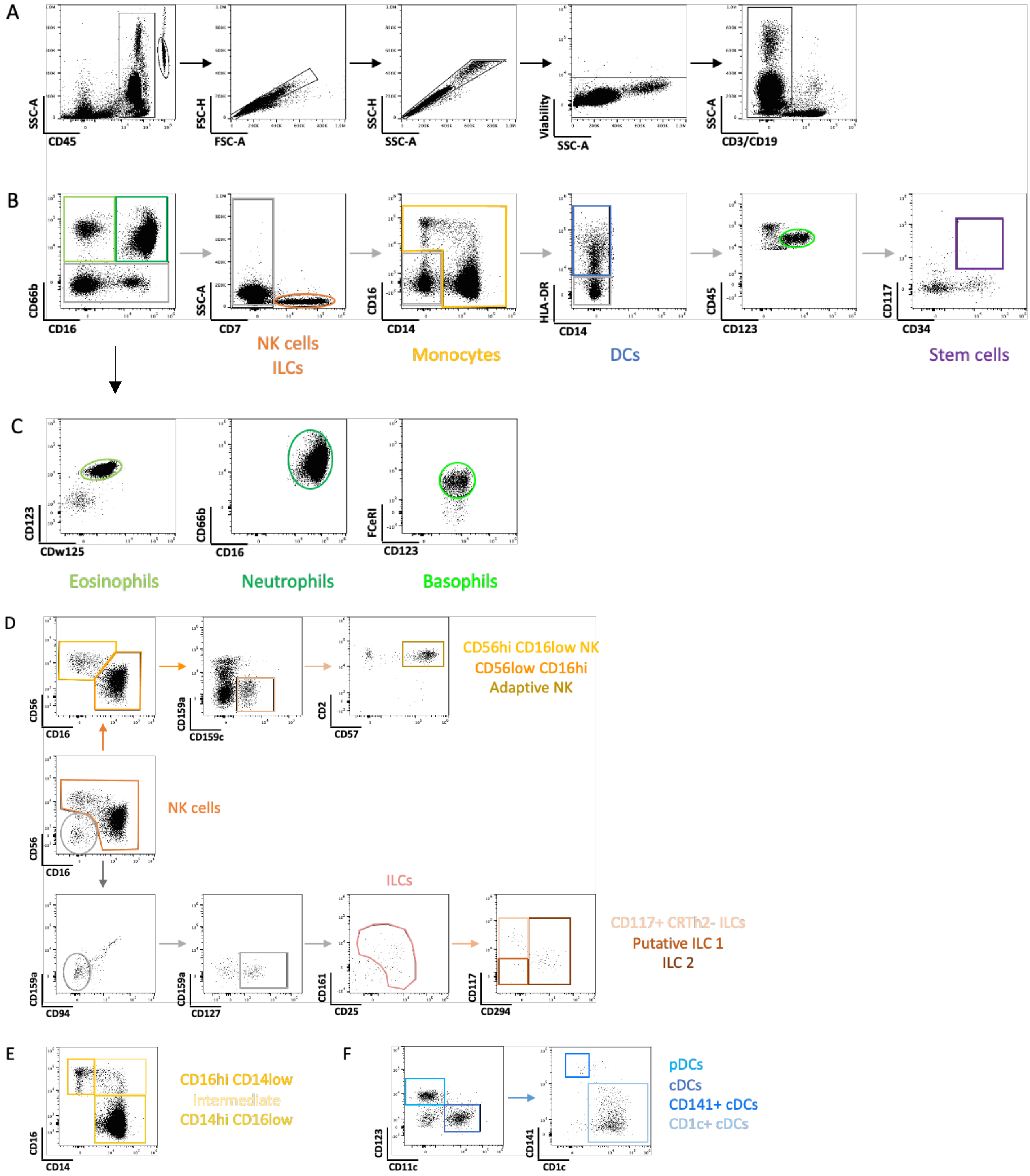


**Supplementary Figure 1.** Gating strategy for innate panel. (A) Innate immune cell populations were identified as live CD45+ singlet cells after the exclusion of the dump channel ($CD3^+$ and $CD19^+$ cells). (B) NK cells, ILCs, monocytes, DCs and stem cells were identified. (C) Polymorphonuclear (PMN) cells were divided into Eosinophils ($CD123^+$ $CDw125^+$), Neutrophils ($CD66b^+$ $CD16^+$) and Basophils ($FCeRI^+$ $CD123^+$). (D) Upon gating on $CD7^+$ cells, NK and ILCs subpopulations could be described. (E) Monocyte subsets were identified with CD16 and CD14 markers. (F) Different DCs subpopulations were segregated upon CD123, CD11c and CD141, CD1c staining. The activation status of PMNs (CD16, CD32, CD63, CXCR4, FCeRI, HLADR, CD62L and PDL1), NK cells (CD56, CD69, CD8a), monocytes (HLA-DR, CD4, PDL-1) and DCs (CD86, CXCR4, CD4, HLA-DR, PDL-1, CD8a) were assessed (not shown).

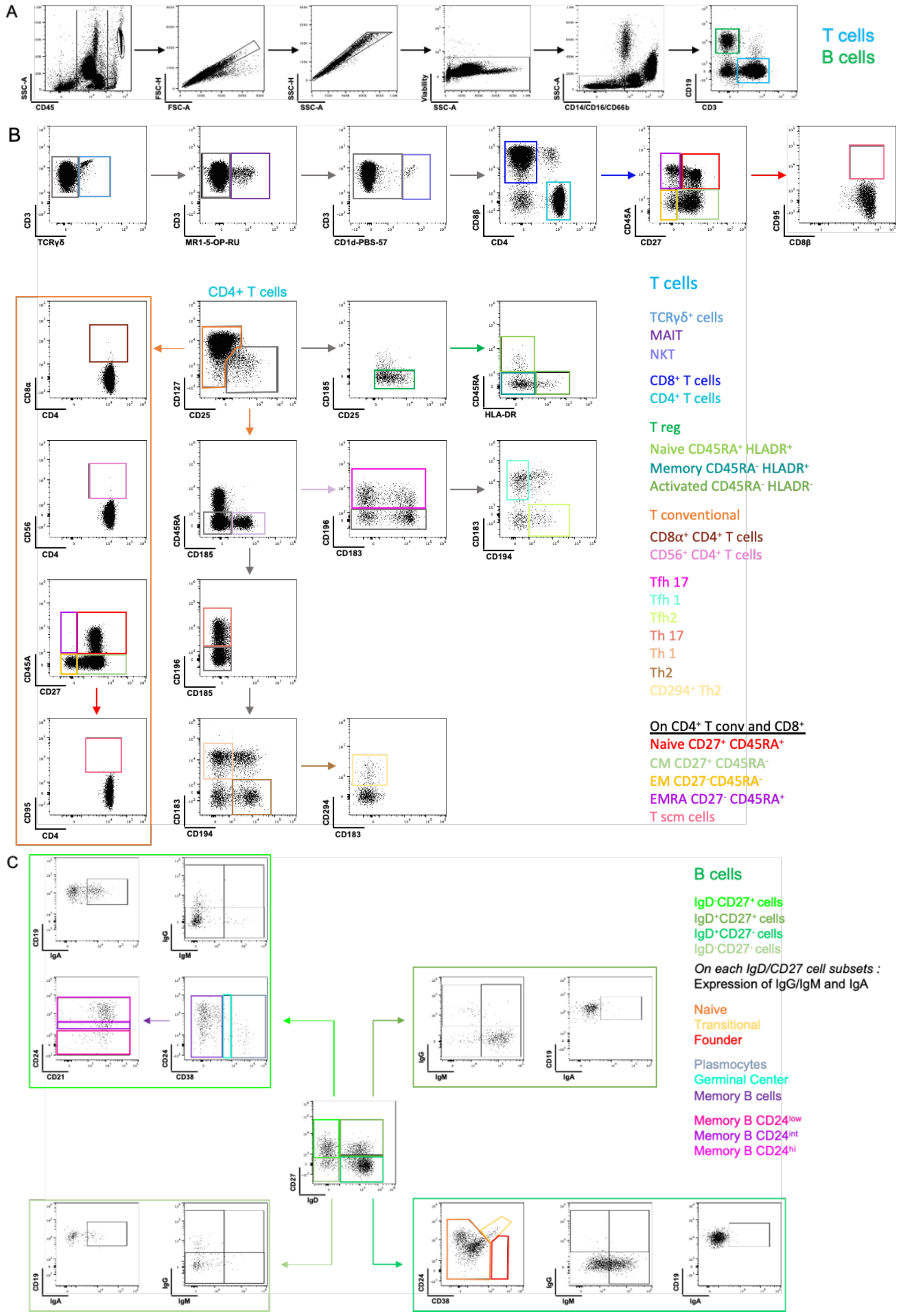


**Supplementary Figure 2.** Gating strategy for adaptive panel**.** Adaptive cells were identified as live CD45$^{+}$ singlet cells exempt from CD14$^{+}$, CD66b$^{+}$ and CD16$^{+}$ cells. T and B cells were characterized as CD3$^{+}$ and CD19$^{+}$, respectively. (B) The major T cell subpopulations such as Tγδ, MAIT, NKT, CD8 and CD4, as well as associated subpopulations (Th and Tfh) were identified among CD3$^{+}$ cells. (C) Within B cells, various markers are used to identify the major B cell sub-populations. The activation status was defined based on the expression of HLA-DR for MAIT and NKT (not shown). The expression of CXCR4, CD8a, CCR7, HLA-DR, PD1, CD95, ICOS and CD56 was used for additional characterization of T cell subsets (not shown).

**INNATE Panel**

| Laser | Specificity | Clone | Fluorophore | Supplier | Cytodelics | CellCover |
|---|---|---|---|---|---|---|
| 355 nm | CDw125 | A14 | BUV395 | BD Biosciences | Red | Red |
| | CD94 | HP-3D9 | BUV496 | BD Biosciences | Green | Red |
| | CD159c | 134591 | BUV563 | BD Biosciences | Red | Red |
| | CD16 | 3G8 | BUV615 | BD Biosciences | Green | Green |
| | CD86 | BU63 | BUV661 | BD Biosciences | Red | Red |
| | CD62L | DREG-56 | BUV737 | BD Biosciences | Red | Red |
| | FcεRI | AER-37 | BUV805 | BD Biosciences | Green | Green |
| 405 nm | CD141 | M80 | BV421 | SONY Biotechnology | Red | Red |
| | CD66b | G10F5 | Pacific blue | SONY Biotechnology | Green | Green |
| | CD159a | 131411 | BV480 | BD Biosciences | Red | Red |
| | CD2 | RPA-2.10 | BV510 | SONY Biotechnology | Green | Green |
| | CD11c | B-ly6 | BV570 | BD Biosciences | Red | Red |
| | CD117 | 104D2 | BV605 | SONY Biotechnology | Red | Red |
| | CD184 | 12G5 | BV650 | BD Biosciences | Red | Red |
| | CD63 | H5C6 | BV711 | SONY Biotechnology | Red | Red |
| | CD56 | 5.1H11 | BV750 | SONY Biotechnology | Green | Red |
| | CD127 | A019D5 | BV785 | SONY Biotechnology | Red | Red |
| 488 nm | CD123 | 6H6 | AF532 | ThermoFisher | Red | Red |
| | CD32 | FLI8.26 | BB630-P2 | BD Biosciences | Red | Red |
| | CD8α | SK1 | PerCP | SONY Biotechnology | Green | Green |
| | CD161 | DX12 | BB700 | BD Biosciences | Red | Red |
| | CD69 | FN50 | PerCP-EF710 | ThermoFisher | Red | Red |
| | CD14 | M5E2 | BB790-P | BD Biosciences | Green | Green |
| 561 nm | CD294 | BM16 | PE | BD Biosciences | Red | Red |
| | CD45 | HI30 | Spark YG™ 593 | SONY Biotechnology | Green | Green |
| | CD25 | BC96 | PE-Dazzle594 | BioLegend | Red | Red |
| | CD7 | M-T701 | PE-Cy5 | BD Biosciences | Red | Red |
| | CD34 | 581 | PE-Cy5.5 | ThermoFisher | Green | Red |
| | CD1c | L161 | PE-Cy7 | SONY Biotechnology | Red | Red |
| 647 nm | CD274 | MIH1 | APC | BD Biosciences | Red | Red |
| | CD57 | HNK-1 | AF647 | SONY Biotechnology | Green | Green |
| | HLA-DR | L243 | AF700 | SONY Biotechnology | Red | Red |
| | CD3 | SK7 | APC-Cy7 | SONY Biotechnology | Red | Red |
| | CD19 | HIB19 | | SONY Biotechnology | Red | Red |
| | CD4 | SK3 | APC/Fire™ 810 | SONY Biotechnology | Green | Green |

**Adaptive Panel**

| Laser | Specificity | Clone | Fluorophore | Supplier | Cytodelics | CellCover |
|---|---|---|---|---|---|---|
| 355 nm | CD183 | 1C6/CXCR3 | BUV395 | BD Biosciences | Red | Red |
| | CD196 | 11A9 | BUV496 | BD Biosciences | Red | Red |
| | CD19 | HIB19 | BUV563 | BD Biosciences | Green | Green |
| | CD21 | B-ly4 | BUV615 | BD Biosciences | Red | Red |
| | CD185 | RF8B2 | BUV661 | BD Biosciences | Red | Red |
| | CD27 | O323 | BUV737 | BD Biosciences | Red | Red |
| | CD3 | UCHT1 | BUV805 | BD Biosciences | Green | Green |
| 405 nm | MR1 5-OP-RU | N/a | BV421 | NIH | Red | Red |
| | IgD | IA6-2 | Pacific Blue | SONY Biotechnology | Red | Red |
| | CD8β | 2ST8.5H7 | BV480 | BD Biosciences | Red | Red |
| | IgM | MHM-88 | BV510 | BioLegend | Red | Red |
| | CD194 | L291H4 | BV605 | SONY Biotechnology | Red | Red |
| | TCRγδ | 11F2 | BV650 | BD Biosciences | Red | Green |
| | IgG | G18-145 | BV711 | BD Biosciences | Red | Red |
| | CD56 | 5.1H11 | BV750 | SONY Biotechnology | Green | Red |
| | CD197 | G043H7 | BV785 | BioLegend | Red | Red |
| 488 nm | IgA | IS11-8E10 | FITC | Miltenyi Biotec | Green | Red |
| | CD45RA | HI100 | AF532 | ThermoFisher | Green | Red |
| | CD8α | SK1 | PerCP | SONY Biotechnology | Green | Green |
| | CD279 | EH12.1 | BB700 | BD Biosciences | Red | Red |
| | CD95 | DX2 | PerCP-EF710 | ThermoFisher | Red | Red |
| 561 nm | CD1d PBS-57 | N/a | PE | NIH | Red | Green |
| | CD45 | HI30 | Spark YG™ 593 | SONY Biotechnology | Green | Green |
| | CD25 | BC96 | PE-Dazzle594 | BioLegend | Red | Red |
| | CD184 | 12G5 | PE-Cy5 | SONY Biotechnology | Red | Red |
| | CD127 | eBioRDR5 | PE-Cy5.5 | ThermoFisher | Green | Green |
| | CD14 | M5E2 | | BD Biosciences | Green | Green |
| | CD66b | G10F5 | PE-Cy7 | ThermoFisher | Green | Green |
| | CD16 | 3G8 | | SONY Biotechnology | Green | Green |
| | CD38 | S17015F | PE/Fire™ 810 | SONY Biotechnology | Green | Red |
| 647 nm | CD278 | C398.4A | APC | SONY Biotechnology | Green | Red |
| | CD294 | BM16 | AF647 | BD Biosciences | Red | Red |
| | HLA-DR | L243 | AF700 | SONY Biotechnology | Red | Red |
| | CD24 | ML5 | APC-Cy7 | BioLegend | Red | Red |
| | CD4 | SK3 | APC/Fire™ 810 | SONY Biotechnology | Green | Green |

**Supplementary Table 2**. Antibody validation for fixed blood and stabilized PBMCs samples. The validation status is color-coded: *Green*: signal comparable to fresh samples; *Red*: no/low/unspecific signal when compared to fresh samples

| | | | MIv3 donor match | |
|---|---|---|---|---|
| **Diagnosis** | **Sex** | **Age** | **Sex** | **Age** |
| sarcoidosis | F | 19 | F | 30 |
| idiopathic hypereosinophilic syndrome | M | 67 | M | 67 |
| mixed connective tissue disease | F | 34 | F | 34 |
| mixed connective tissue disease | F | 34 | F | 34 |
| giant cell arteritis | F | 76 | F | 76 |
| ANCA vasculitis | F | 71 | F | 71 |
| uveitis | F | 72 | F | 69 |
| systemic lupus erythematosus | F | 26 | F | 30 |

**Supplementary Table 3.** Patient metadata

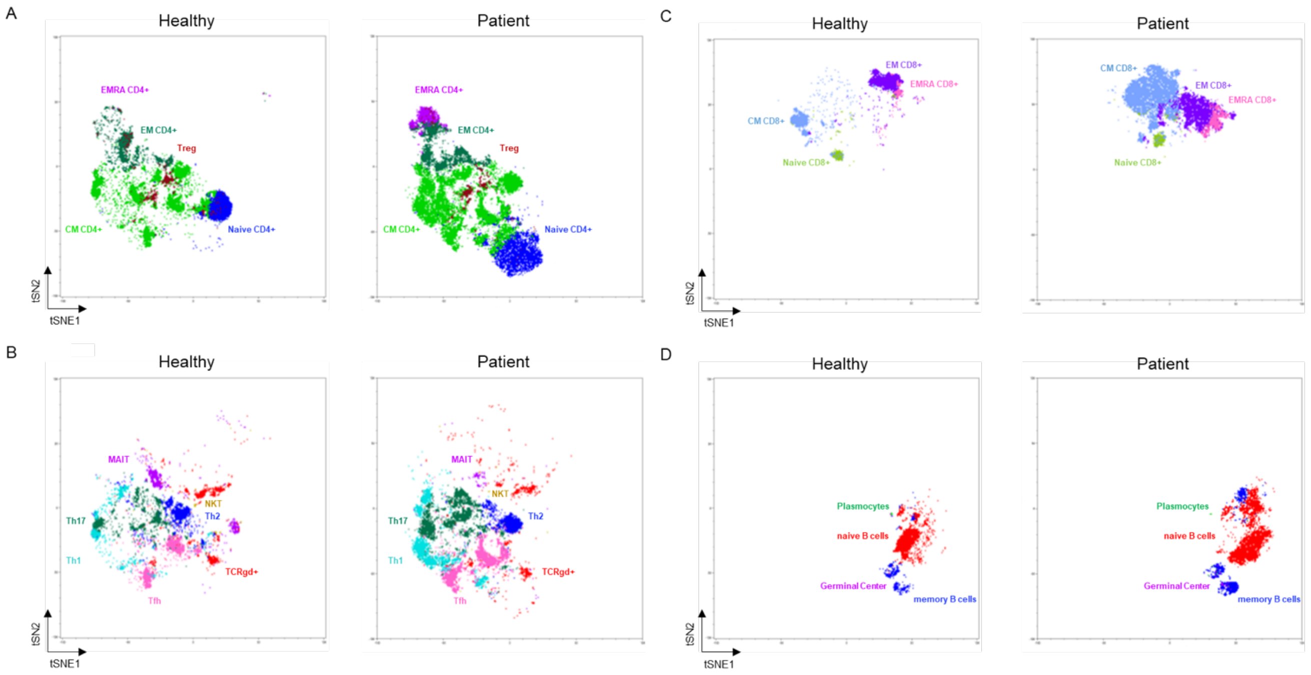


**Supplementary Figure 3**. Non-supervised analysis of adaptive cell subsets. Non-supervised analysis was performed by Sony prototype software. tSNE projections of clusters were annotated based on manual gating. A representative example of a healthy donor (left) and of a diseased donor (right) is shown. A) $CD4^{+}$ T cells (naïve, CM, EM, EMRA, Treg); B) CD4 subsets (Th, MAIT, NKT, Tγδ); C) $CD8^{+}$ T cells (naïve, CM, EM, EMRA) and D) B cells (plasmocytes, GC, Memory, naïve).
